# Hyperactive TORC1 sensitizes yeast cells to endoplasmic reticulum stress by compromising cell wall integrity

**DOI:** 10.1101/605824

**Authors:** Khadija Ahmed, David E. Carter, Patrick Lajoie

## Abstract

The disruption of protein folding homeostasis in the endoplasmic reticulum (ER) results in an accumulation of toxic misfolded proteins and activates a network of signaling events collectively known as the unfolded protein response (UPR). While UPR activation upon ER stress is well characterized, how other signaling pathways integrate into the ER proteostasis network is unclear. Here, we sought to investigate how the target of rapamycin complex 1 (TORC1) signaling cascade acts in parallel with the UPR to regulate ER stress sensitivity. Using *S. cerevisiae*, we found that TORC1 signaling is attenuated during ER stress and constitutive activation of TORC1 increases sensitivity to ER stressors such as tunicamycin and inositol deprivation. This phenotype is independent of the UPR. Transcriptome analysis revealed that TORC1 hyperactivation results in cell wall remodelling. Conversely, hyperactive TORC1 sensitizes cells to cell wall stressors, including the antifungal caspofungin. Elucidating the crosstalk between the UPR, cell wall integrity, and TORC1 signaling may uncover new paradigms through which the response to protein misfolding is regulated, and thus have crucial implications for the development of novel therapeutics against pathogenic fungal infections.

**IMPORTANCE:** The prevalence of pathogenic fungal infections, coupled with the emergence of new fungal pathogens, has brought these diseases to the forefront of global health problems. While antifungal treatments have advanced over the last decade, patient outcomes have not substantially improved. These shortcomings are largely attributed to the evolutionary similarity between fungi and humans, which limits the scope of drug development. As such, there is a pressing need to understand the unique cellular mechanisms that govern fungal viability. Given that *Saccharomyces cerevisiae* is evolutionarily related to a number of pathogenic fungi, and in particular to the *Candida* species, most genes from *S. cerevisiae* are highly conserved in pathogenic fungal strains. Here we show that hyperactivation of TORC1 signaling sensitizes *S. cerevisiae* cells to both endoplasmic reticulum stress and cell wall stressors by compromising cell wall integrity. Therefore, targeting TORC1 signaling and endoplasmic reticulum stress pathways may be useful in developing novel targets for antifungal drugs.

## INTRODUCTION

The ability of cells to respond to detrimental stresses, such as an aberrant accumulation of toxic misfolded proteins, dictates cell fate under both normal and pathological conditions. Loss of secretory protein homeostasis due to pharmacological, genetic, or environmental perturbations activates a plethora of adaptive responses to help cells overcome the stress (1, 2). In yeast, the ER resident protein Ire1 detects changes in the ER misfolded protein and activates a transcriptional response termed the unfolded protein response (UPR; (3–7). Upon induction of ER stress, the ER chaperone, Kar2, dissociates from the luminal domain of Ire1, allowing it to oligomerize, trans-autophosphorylate, and subsequently activate its cytosolic RNase activity (4, 5, 8–10). Ire1 then splices *HAC1* mRNA to generate a functional variant of the transcript, which upon translation functions as a transcription factor to upregulate genes involved in ER quality control machinery and ribosome biogenesis (5, 8). Cellular adaptation to ER stress is not only dependent on the amplitude of the UPR signal, but also on the selective expression of UPR target genes capable of overcoming a particular stress condition (11). Interestingly, Pincus *et al.* (2014) show that *S. cerevisiae* amplify the UPR with time delayed Ras/PKA signaling, indicating that the response to ER stress is not limited to the UPR (12). Moreover, induction of ER stress activates transcription of genes associated with other types of stress responses (2). Therefore, elucidating how the UPR integrates with other signaling pathways under conditions of ER stress is essential to understand how proteostasis is mediated in the cell.

Given that protein folding in the ER is a highly energetically demanding process, low nutrient status is a potent trigger of the UPR (13). Therefore, the interconnection between metabolic regulation and the UPR is a crucial area of study, one that has thus far been inadequately addressed. Accumulating evidence suggests that the cellular metabolism mediating AMPK signaling cascade and its subsequent regulation of crucial proteins acetyl-CoA carboxylase and TOR, may cooperate with the UPR to mediate cell viability under conditions of ER stress (13–15); however, the mechanisms behind this crosstalk remain to be elucidated. In yeast, TORC1 inhibition with rapamycin protects yeast cells from ER stress-induced vacuolar fragmentation and promotes antifungal synergism (16). In addition, pharmalogical induction of ER stress triggers autophagy, a process negatively regulated by TORC1 (17). It therefore appears that TOR signaling is an important determinant of the yeast ER stress response.

In *S. cerevisiae*, TOR kinases are evolutionarily conserved serine/threonine kinases that function at the core of signaling networks involved in cell growth, metabolism, and nutrient and hormone sensing (18, 19). These TOR kinases are the central component of two distinct complexes: TOR complex 1 (TORC1) and TOR complex 2 (TORC2), of which only TORC1 is rapamycin sensitive (20). In particular, the TORC1 signaling network mediates anabolism and catabolism by coordinating cellular and metabolic processes such as transcription, protein translation, ribosome biogenesis, and cellular architecture (20–23). In addition to mediating anabolic processes, TORC1 promotes cell growth by inhibiting a number of stress response pathways (21, 24, 25). Nevertheless, the manner in which the secretory and TORC1 signaling pathway act in parallel, under conditions of ER stress, remains to be elucidated.

To study the effect of TORC1 signaling on protein folding homeostasis, we employed a hyperactive variant of the TOR1 kinase (*TOR1*^*L2134M*^) and assessed yeast sensitivity to ER stress. We elucidate a novel interplay between proteostasis and TORC1 signaling and show that attenuation of TORC1 signaling is required for adaptation to ER stress. On the other hand, constitutive activation of TORC1 confers increased sensitivity to ER stressors, including the antifungal caspofungin, by compromising cell wall architecture. Our study, therefore, expands the role of ER homeostasis beyond the UPR and defines how TORC1 signaling contributes to the ER stress response.

## RESULTS AND DISCUSSION

### Hyperactive *TOR1*^*L2134M*^ sensitizes cells to ER stress

Previous studies show that the TOR pathway links nutrient status to cell growth and ribosome biogenesis, under conditions of protein misfolding stress (26–28). However, it remains unclear to what extent modulation of TORC1 signaling is required for adaptation to ER stress. Thus, we sought to investigate the effects of TORC1 signaling on the sensitivity to ER stress.

The phosphorylation of the ribosomal protein, Rps6, is regulated in a TORC1-dependent manner and serves as a valid readout for TORC1 activity *in vivo* (29, 30). Previous reports indicate that under conditions of oxidative- and proteotoxic stress, RPS6 phosphorylation is dramatically reduced (31, 32). Therefore, we sought to investigate whether ER stress reduces Rps6 phosphorylation in cells with hyperactive TORC1 signaling (Fig. 1A). As such, cells expressing either WT *TOR1* or hyperactive *TOR1*^*L2134M*^ were treated with the canonical ER stress inducer, tunicamycin (Tm; Fig. 1B). Tm is a potent inducer of the UPR as it inhibits N-glycosylation of proteins, prevents proper protein folding, and thereby causes an accumulation of misfolded proteins in the ER (33). While the addition of Tm (2.5 ug/mL) significantly decreased Rps6 phosphorylation in cells expressing WT *TOR1*, there was no significant difference in cells expressing hyperactive *TOR1*^*L2134M*^ (Fig. 1B-C). Rapamycin, an inhibitor of TORC1, was used as a positive control, for Sch9 downregulation. Combined with previous studies showing that phosphorylation of Sch9, another TORC1 effector, is decreased during Tm treatment (34), our results suggest that TORC1 deactivation plays an important role in ER stress tolerance. As such, we then sought to determine how impacting proper TORC1 signaling affects the cell’s response to ER stressors.

**Figure 1:**
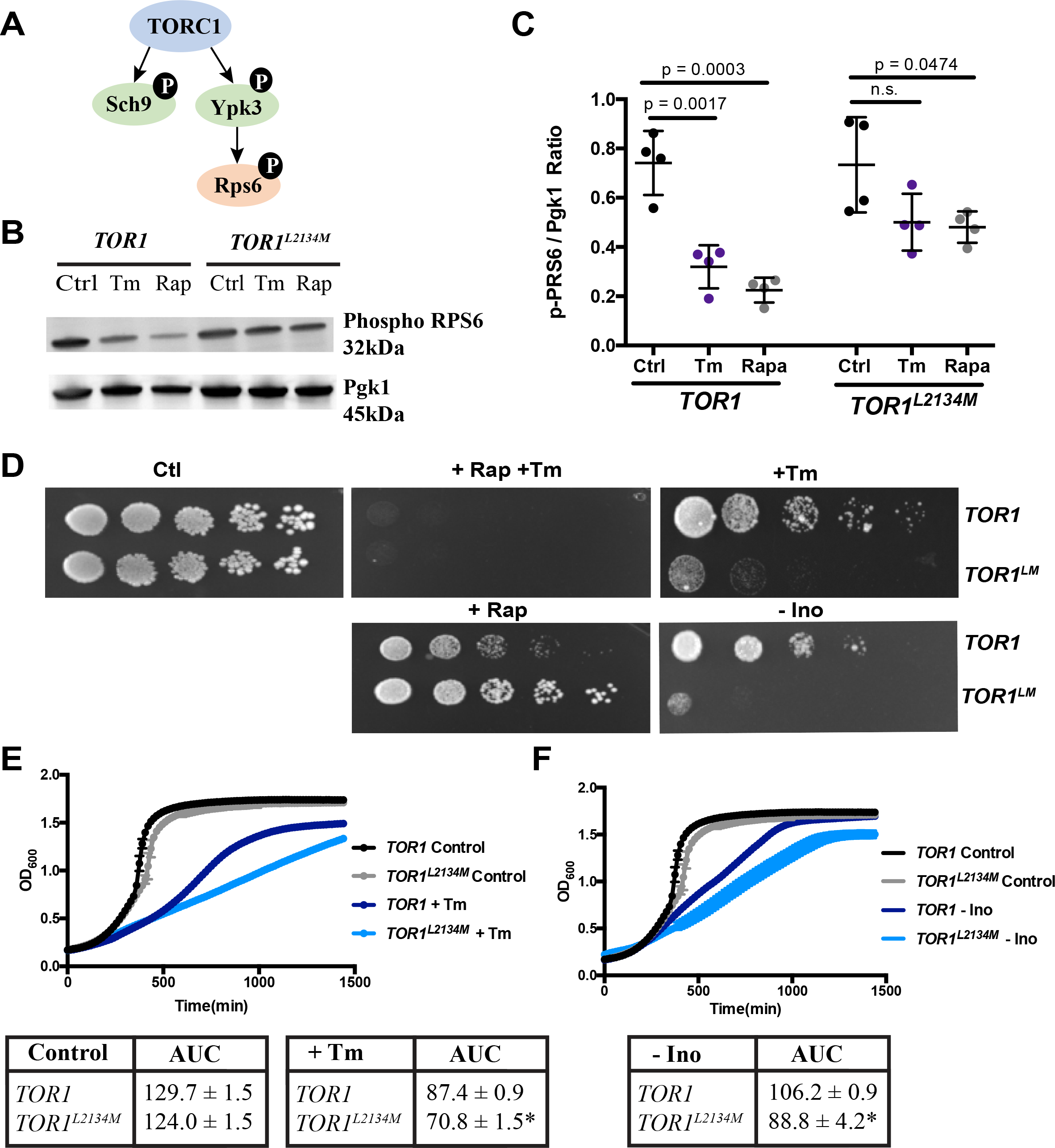
Cells expressing hyperactive *TOR1*^*L2134M*^ are more sensitive to ER stress. **(A)** Representative schematic of the downstream targets of TORC1 kinase activity. **(B)** Western blot analysis of Rps6 phosphorylation following treatment with tunicamycin (Tm; 2.5 μg/mL) or rapamycin (Rap; 200 ng/mL). Pgk1 was used as a loading control. **(C)** Quantification of (B). Rps6 phosphorylation is not significantly attenuated in hyperactive *TOR1*^*L2134M*^ cells following treatment with tunicamycin (n=4; ± SD). **(D)** Cell growth of WT *TOR1* and *TOR1*^*L2134M*^ cells was assessed by serial dilutions on YPD plates supplemented with rapamycin (Rap; 10 ng/mL), tunicamycin (Tm; 1.0 μg/mL), both Rap and Tm, or SC plates supplemented without inositol (+/− Inositol). Cells expressing hyperactive *TOR1*^*L2134M*^ were more resistant to rapamycin treatment and more sensitive to tunicamycin stress and inositol withdrawal. **(E-F)** Liquid growth assays of yeast cells expressing WT *TOR1* and *TOR1*^*L2134M*^ were used to further assess sensitivity to tunicamycin stress (Tm; 1.0 μg/mL) and inositol withdrawal (-Ino). Data is quantified as area under the curve (AUC; *p < 0.01; mean ± SD; n=3). All conditions were run simultaneously. Control conditions are reproduced on both panels for clarity.

First, we assessed cell growth in the presence of both Tm and the TORC1 inhibitor, rapamycin (Fig. 1D). We found that rapamycin treatment exacerbates the growth defect caused by Tm-induced ER stress (Fig. 1D). Similarly, cells expressing a rapamycin-resistant hyperactive *TOR1*^*L2134M*^ (24) displayed an increased growth defect upon Tm stress (Fig. 1D). To investigate the effects of hyperactive *TOR1* on a more physiologically relevant ER stressor, cells were exposed to conditions of inositol withdrawal. While it is unclear how exactly inositol deprivation triggers UPR activation, some studies have postulated that it triggers the UPR by either changing the lipid composition of the ER membrane (35–37) or by impairing membrane trafficking (38, 39). In contrast to cells expressing WT *TOR1*, cells expressing the hyperactive allele were inositol auxotrophs (Fig. 1D). Increased ER stress sensitivity of *TOR1*^*L2134M*^ was confirmed using liquid growth assays (Fig. 1E-F). As expected, compared to cells expressing WT *TOR1,* cells expressing hyperactive *TOR1*^*L2134M*^ had a significant growth defect following treatment with Tm (Fig. 1E) or inositol withdrawal (Fig. 1F). Interestingly, we previously showed that TOCR1 hyperactivation using the *TOR1*^*L2134M*^ strain also sensitizes yeast to expanded polyglutamine proteins (40) linked to ER stress and UPR activation in yeast and other models of Huntington’s disease (41, 42). Taken together, our results indicate that defective TORC1 signaling increases sensitivity to canonical ER stressors. Both phenotypes can be linked to a defective response to ER stress.

### Cells expressing hyperactive *TOR1*^*L2134M*^ have a functional UPR

Having shown that cells expressing hyperactive *TOR1*^*L2134M*^ are more sensitive to ER stress, we next sought to examine whether this increased sensitivity was due to defects in the ability to activate the UPR. As previously described, under conditions of ER stress, the ER protein folding sensor, Ire1, splices *HAC1* mRNA to produce an active transcription factor (4). We therefore assessed the ability of Ire1 to splice *HAC1* mRNA using RT-PCR (Fig. 2A-B). Surprisingly, inositol withdrawal induced *HAC1* splicing in both WT *TOR1* and hyperactive *TOR1*^*L2134M*^ mutants (Fig. 2A, arrow). Additionally, after 1 hr of treatment with Tm, cells expressing hyperactive *TOR1*^*L2134M*^ spliced *HAC1* mRNA, and this response was still evident after 2 hrs of induction, as indicated by a smaller fragment in the agarose gel (Fig. 2B, arrow). As a whole, these results indicate that increased ER sensitivity of cells expressing hyperactive *TOR1*^*L2134M*^ is not due to impaired functionality of the UPR.

**Figure 2:**
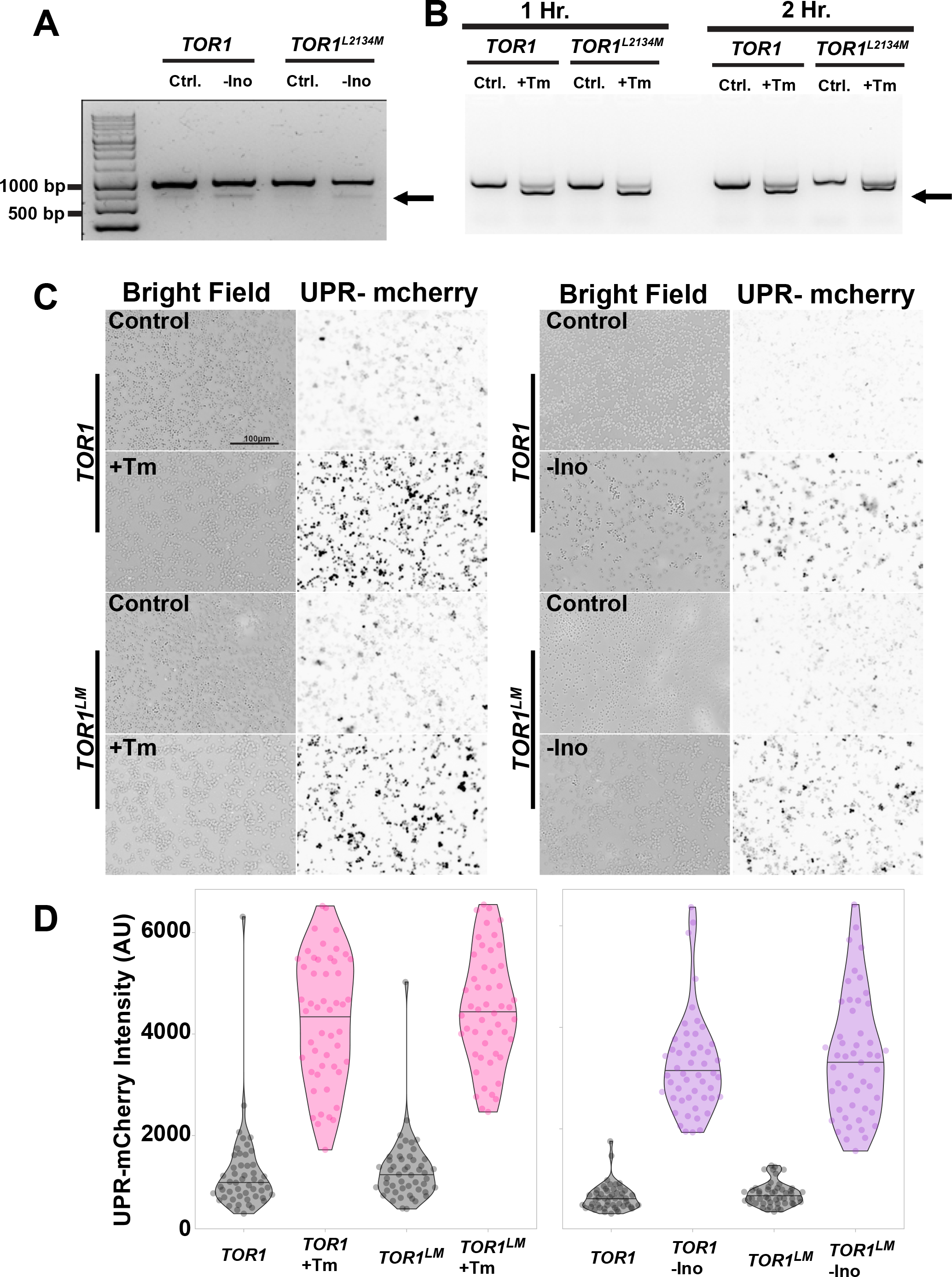
The UPR is not impaired in yeast cells expressing hyperactive *TOR1*^*L2134M*^. **(A)** Treatment with ER stressors induces *HAC1* mRNA splicing. WT *TOR1* and hyperactive *TOR1*^*L2134M*^ mutant were either untreated (Ctrl.), subjected to inositol withdrawal (-Ino) for 2hrs, or **(B)** treated with tunicamycin (Tm; 1.0 μg/mL) for up to 2 hrs. RT-PCR was conducted using *HAC1* primers. Arrows indicate Ire1 mediated *HAC1* splicing. **(C)** Representative fluorescence microscopy images of WT *TOR1* and *TOR1*^*L2134M*^ cells expressing UPR-mcherry fluorescent reporters, following treatment with tunicamycin (Tm; 1.0 μg/mL) and inositol withdrawal (-Ino) for 2 hours. **(D)** Quantification of (C).

Spliced *HAC1* mRNA is translated into an active transcription factor, which then translocates to the nucleus where it binds to unfolded protein response element (UPRE) sequences in gene promoters^44^. In response to ER stress, Hac1 alone activates over 400 UPR target genes, including ER chaperones, genes that mediate membrane expansion, and genes involved in ribosome biogenesis (1, 43, 44). As such, increased sensitivity to ER stress may be due to an inability to transcriptionally activate the UPR. We tested this possibility by transforming a UPRE-mcherry fluorescent reporter (45) into cells expressing *TOR1* and *TOR1*^*L2134M*^ and assessing UPR activation with fluorescence microscopy (Fig. 2C-D). Surprisingly, there was no significant difference between cells expressing *TOR1* and hyperactive *TOR1*^*L2134M*^ in their ability to activate the UPR under conditions of Tm stress and inositol withdrawal. Additionally, we quantitatively assessed the mRNA levels of the yeast resident chaperone and canonical UPR target gene, *KAR2*, using qRT-PCR (Fig. 3A). In line with our previous data, hyperactive *TOR1*^*L2134M*^ was able to increase the expression of *KAR2*, following treatment with Tm and inositol withdrawal. Taken together, these results suggest that the increased sensitivity of cells expressing *TOR1*^*L2134M*^ to ER stress is unlikely to be due to impaired UPR activation.

**Figure 3:**
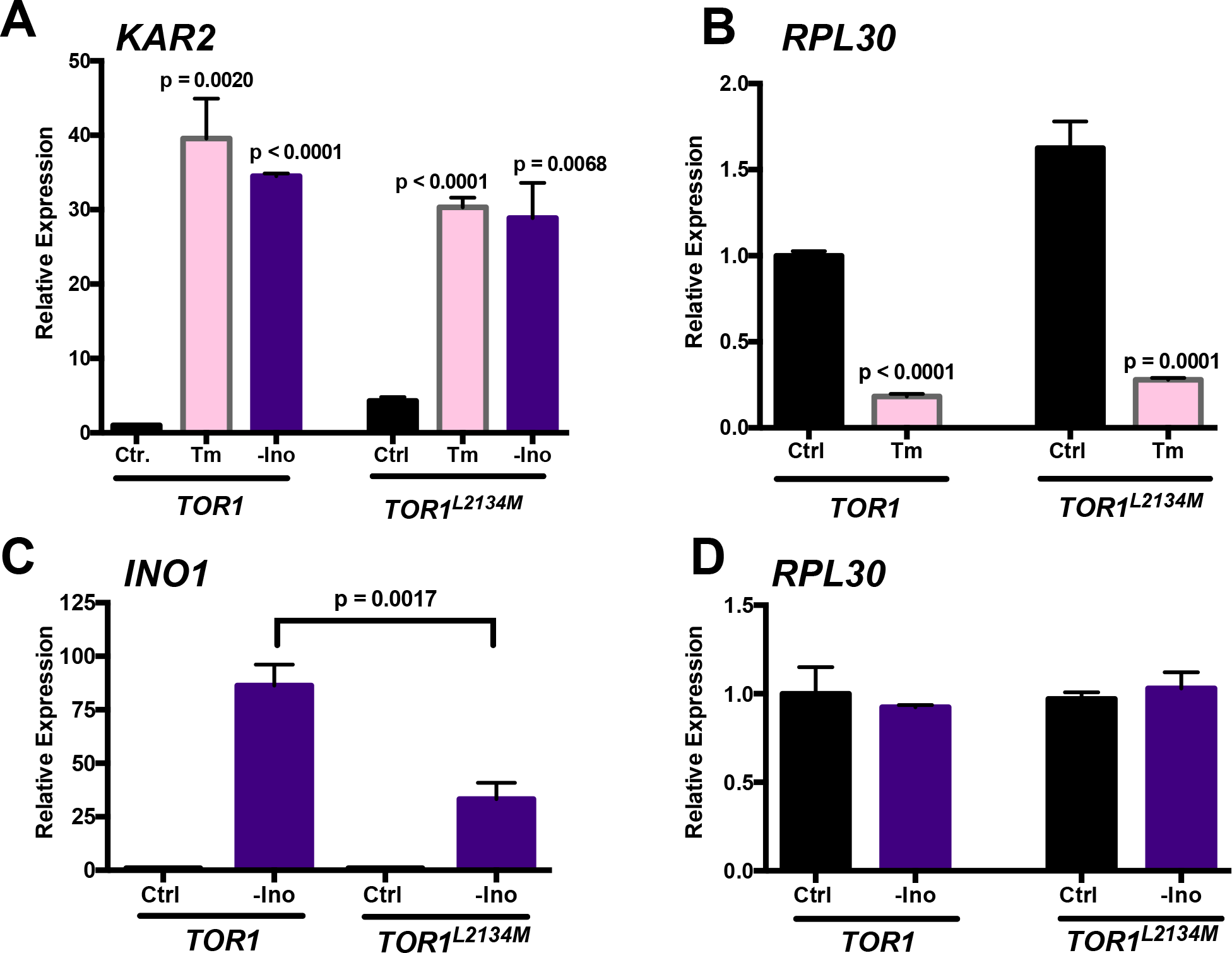
Hyperactive *TOR1*^*L2134M*^ can transcriptionally activate the UPR, but has impaired inositol synthesis. **(A)** Hyperactive *TOR1*^*L2134M*^ can upregulate expression of the ER chaperone *KAR2* following treatment with tunicamycin (Tm; 2.5 μg/mL) or inositol withdrawal (-Ino) (n =3; ± SD). **(B)** Following treatment with tunicamycin stress (Tm; 2.5 μg/mL), hyperactive *TOR1*^*L2134M*^ can downregulate expression of *RPL30* (n=3; ± SD). **(C)** Under conditions of inositol withdrawal (-Ino), cells expressing hyperactive *TOR1*^*L2134M*^ have impaired synthesis of *INO1*. Cells expressing WT *TOR1* and hyperactive *TOR1*^*L2134M*^ were treated with inositol withdrawal for 2 hrs. qRT-PCR was conducted using *INO1* primers (n= 3; ± SD). **(D)** Inositol withdrawal does not induce downregulation of *RPL30*. Cells expressing WT *TOR1* and hyperactive *TOR1*^*L2134M*^ were subjected to inositol withdrawal for 2 hrs (n=3; ± SD).

Additionally, actively dividing yeast allocate up to 85% of their transcriptional activity to ribosome biogenesis (46); however, under conditions of ER stress, there is a downregulation in the expression of ribosome genes in order to increase the expression of UPR target genes (47, 48). As such, we employed qRT-PCR to assess the expression of *RPL30*, a gene involved in ribosome biogenesis (Fig. 3B). Cells expressing hyperactive *TOR1*^*L2134M*^ significantly downregulated expression of *RPL30* (Fig. 3B). This is probably due to the fact that multiple pathways regulate ribosome biogenesis. For example, PKA deactivation during ER stress is also responsible for repressing transcription of ribosomal protein genes (12). Furthermore, depleting inositol triggers the ER stressor, Ire1, which induces transcription of the inositol biosynthetic gene, *INO1* (8, 49). Therefore, we investigated whether the inositol auxotrophy of cells expressing *TOR1*^*L2134M*^ was due to the inability to synthesize *INO1*. Cells expressing *TOR1* and *TOR1*^*L2134M*^ were treated with inositol withdrawal and qRT-PCR was conducted to assess the expression of *INO1* and *RPL30* (Fig. 3C-D). Interestingly, hyperactive *TOR1*^*L2134M*^ impaired the transcription of *INO1* (Fig. 3C) but did not impair ribosome biogenesis (Fig. 3D). Taken together, these results suggest that under conditions of ER stress, cells expressing hyperactive *TOR1*^L2134M^ are defective in regulating *INO1* transcription.

### Defects in cell wall integrity underlie *TOR1*^*L2134M*^ sensitivity to ER stress

Despite having a functional UPR, our studies show that cells expressing hyperactive *TOR1*^*L2134M*^ have increased sensitivity to canonical ER stressors. Therefore, to assess how ER stress alters the transcriptome in hyperactive *TOR1*^*L2134M*^ mutants, we treated two independent cultures of WT *TOR1* and hyperactive *TOR1*^*L2134M*^ cells with Tm and used microarray analysis to uncover genes that were differentially expressed in hyperactive *TOR1*^*L2134M*^ cells (Fig. 4A-D). Data was analyzed by filtering for genes that showed a two-fold change in expression with a p value < 0.05. The transcripts of the genes that were differentially downregulated (Fig. 4C) and upregulated (Fig. 4D) were categorized based on their cellular components using the yeast SGD GO term finder. Interestingly, among the genes that were upregulated, a large majority encoded proteins that localized to the cell periphery and plasma membrane (Fig. 4D). Of note, genes encoding three cell wall incorporated mannoproteins, *FIT1, FIT2*, and *FIT3* were upregulated in hyperactive *TOR1*^*L2134M*^ cells (Fig. 4D). Fit proteins are involved in iron uptake (50). Validation with qRT-PCR revealed that hyperactive *TOR1*^*L2134M*^ cells had significantly higher steady-state levels of *FIT1, FIT2*, and *FIT3*, compared to cells expressing WT *TOR1* (Fig. 4E-G). Interestingly, *FIT* genes are also upregulated in cells carrying deletions in genes encoding the phosphatases *PTC1* and *PTC6* that displayed compromised TORC1 signaling (51). Additionally, the expression of both *FIT2* and *FIT3* was significantly higher compared to WT *TOR1* cells following treatment with Tm (Fig. 4F-G). Interestingly, increased mannoprotein levels is observed in cells with compromised cell wall (52). Taken together, these results suggest that hyperactive *TOR1*^*L2134M*^ alters the cell wall composition of yeast cells.

**Figure 4:**
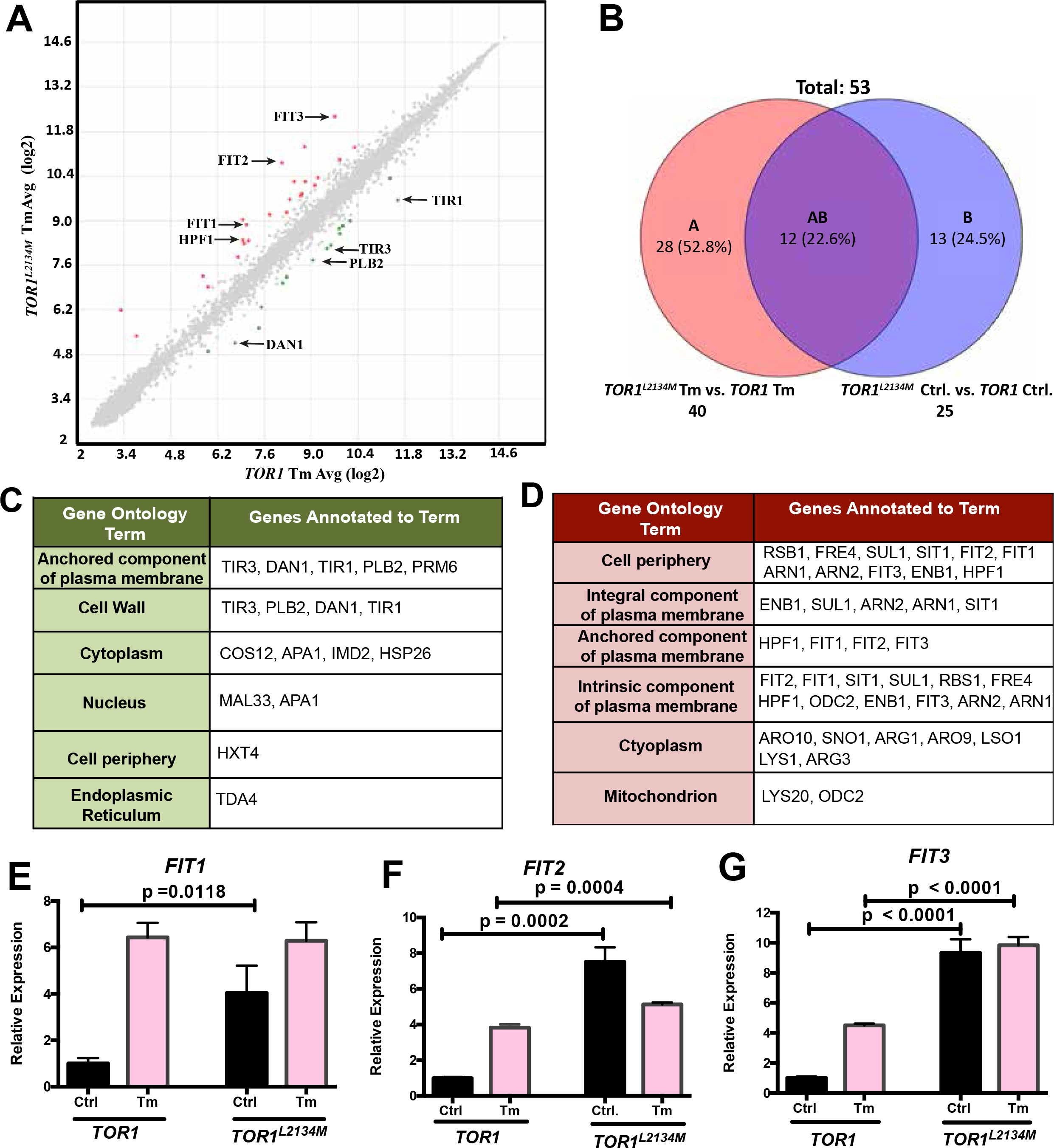
ER stress induces a change in the cell wall composition of cells expressing hyperactive *TOR1*^*L2134M*^. **(A)** Microarray analysis of genes differentially expressed in yeast cells expressing WT *TOR1* or hyperactive *TOR1*^*L2134M*^, following treatment with tunicamycin (Tm; 2.5 μg/mL). Arrows indicate cell wall genes that are differentially expressed in cells expressing hyperactive *TOR1*^*L2134M*^. **(B)** Microarray analysis of genes differentially expressed in *TOR1* and *TOR1*^*L2134M*^ control cells compared to *TOR1* and *TOR1*^*L2134M*^ cells treated with tunicamycin (Tm; 2.5 μg/mL). **(C)** Genes downregulated two-fold in hyperactive *TOR1*^*L2134M*^ cells in response to tunicamycin stress (Tm; 2.5 μg/mL). **(D)** Genes upregulated two-fold in hyperactive *TOR1*^*L2134M*^ cells in response to tunicamycin stress. Gene ontology lists were generated with the gene ontology term finder on the *Saccharomyces* genome database. Numerous cell wall genes are differentially expressed in hyperactive *TOR1*^*L2134M*^ cells compared to cells expressing WT *TOR1*. **(E)** qRT-PCR was used to validate the microarray analysis and assess expression of mannoprotein genes *FIT1*, **(F)** *FIT2*, and **(G)** *FIT3* following treatment with tunicamycin (Tm; 2.5 μg/mL; n=3; ± SD).

ER stress tolerance in yeast depends on the activation of the cell wall integrity pathway, which is, in part, regulated by TORC1 (53–57). Additionally, cells with defects in cell wall integrity exhibit inositol auxotrophy (58). As such, we investigated whether the increased sensitivity of cells expressing hyperactive *TOR1*^*L2134M*^ was due to defects in cell wall integrity. A general approach to assess whether a specific phenotype is due to a cell wall defect is to test the remediating effects of the cell wall stabilizer sorbitol (59). Interestingly, supplementing with sorbitol rescued the toxicity caused by Tm stress in hyperactive *TOR1*^*L2134M*^ mutants (Fig. 5A), suggesting that these cells have a defective cell wall. To further examine cell wall composition, cells expressing *TOR1* and *TOR1*^*L2134M*^ were treated with the cell wall antagonist, calcofluor white (CFW) and liquid growth assays were assessed (Fig. 5B). In line with our previous results, cells expressing hyperactive *TOR1*^*L2134M*^ were significantly more sensitive to CFW than cells expressing WT *TOR1* (Fig. 5B). Previous literature indicates that due to increased activation of cell wall stress responses, yeast strains with defects in cell wall integrity have a greater deposition of chitin in their cell wall and become more sensitive to the CFW (60). Therefore, cells expressing *TOR1* and *TOR1*^*L2134M*^ were stained with CFW and chitin staining was analyzed using fluorescence microscopy and flow cytometry (Fig. 5C). Compared to WT *TOR1* cells, cells expressing hyperactive *TOR1*^*L2134M*^ appeared more clustered and displayed significantly more chitin content (Fig. 5C). Taken together, our data suggests that the increased sensitivity of hyperactive *TOR1*^*L2134M*^ mutants can be traced back to defects in cell wall integrity.

**Figure 5:**
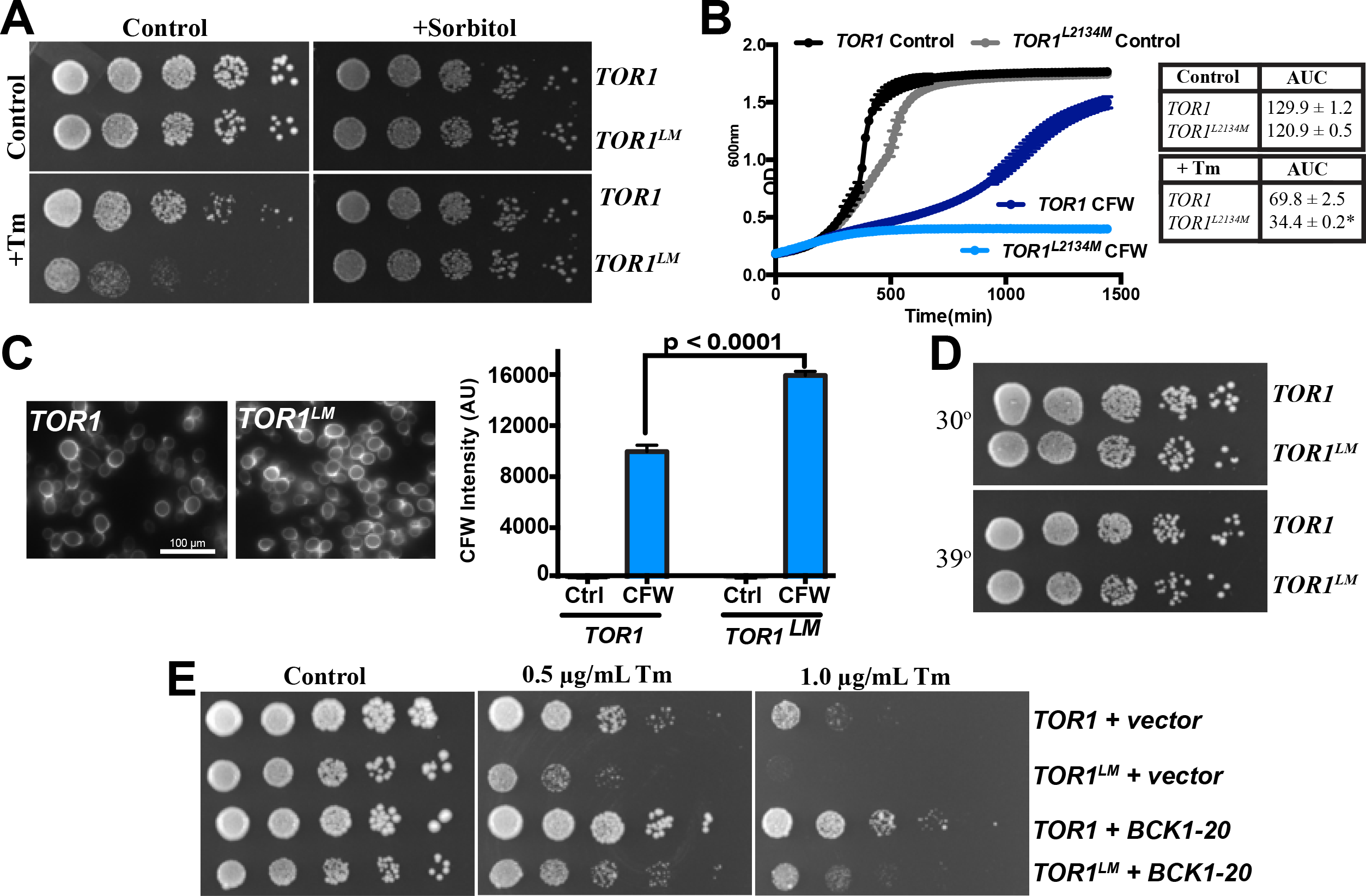
Increased sensitivity of hyperactive *TOR1*^*L2134M*^, in response to ER stress, is due to defects in cell wall integrity. **(A)** Cell growth of WT *TOR1* and *TOR1*^*L2134M*^ cells was assessed by serial dilutions on YPD plates supplemented with various concentrations of tunicamycin (Tm), sorbitol (1 M), or both tunicamycin and sorbitol. Sorbitol rescues tunicamycin toxicity caused by hyperactive *TOR1*^*L2134M*^. **(B)** Liquid growth assay of *TOR1* and *TOR1*^*L2134M*^ cells following treatment with calcofluor white (CFW; 20 μg/mL). Data was quantified by measuring area under the curve (AUC; n=3; *p < 0.001; mean ± SD). **C)** Representative fluorescence microscopy images of cells expressing WT *TOR1* and hyperactive *TOR1*^*L2134M*^, following treatment with calcofluor white (CFW; 20 μg/mL). Cells expressing hyperactive *TOR1*^*L2134M*^ are aggregated and have increased fluorescence, corresponding to an increase in chitin synthesis (Left panel). Flow cytometric analysis of cells treated with calcofluor white (CFW; 2.5 μg/mL). Cells expressing hyperactive *TOR1*^*L2134M*^ have significantly higher mean fluorescence intensity compared to WT TOR cells (right panel; n = 3; mean ± SD). **(D)** Growth of WT *TOR1* and *TOR1*^*L2134M*^ cells in response to elevated temperature was assessed by serial dilution on YPD plates. There was no growth defect caused by hyperactive *TOR1*^*L2134M*^. **(E)** Cell growth of WT *TOR1* and *TOR1*^*L2134M*^ transformed with either an empty vector or BCK1-20 was assessed by serial dilution on SC-ura plates supplemented with various concentrations of tunicamycin (Tm).

Consistent with a defect in cell wall biogenesis, loss of function of any kinase downstream of the canonical MAPK cell wall integrity pathway (CWI) results in growth defects at elevated temperatures (61–64). Therefore, we investigated whether the increased sensitivity of hyperactive *TOR1*^*L2134M*^ to ER stress could be attributed to defects in the canonical CWI pathway. Surprisingly, compared to WT *TOR1* cells, cells expressing hyperactive *TOR1*^*L2134M*^ showed no growth defect at elevated temperatures (Fig. 5D). To further investigate whether the CWI pathway was impaired, we assessed the effects of constitutive activation of the CWI pathway by transforming a hyperactive *BCK1-20* allele into WT *TOR1* and hyperactive *TOR1*^*L2134M*^ cells (Fig. 5E). Interestingly, *BCK1-20* overexpression equally rescued Tm toxicity in both WT *TOR1* and hyperactive *TOR1*^*L2134M*^ cells (Fig. 5E), with *TOR1*^*L2134M*^ cells still displaying increased sensitivity compared to wild-type. These results indicate that other regulators of the cell wall composition downstream of Bck1 may be defective in the mutant cells.

### Hyperactive *TOR1*^*L2134M*^ cells have defects in glucan synthase expression and are more sensitive to caspofungin

Within the host organism, pathogenic fungi face numerous environmental stressors such as low nutrient availability and changes in pH and temperature (65, 66). As such, the fungal cell wall acts as the first line of defense, providing a rigid cellular boundary to withstand internal turgor pressure and extracellular stresses (67). Proper cell wall architecture requires three major components: β-1-3-glucan, chitin, and mannoproteins– all of which come together to form a large macromolecular complex (67, 68). Our results indicate that cells expressing hyperactive *TOR1*^*L2134M*^ increase expression of mannoprotein genes as well as chitin aggregation, both of which are phenotypes associated with impaired β-1-3-glucan synthesis (69–71). To test this possibility, we used qRT-PCR to assess the expression of the β-1-3-glucan synthase genes, *FKS2* and *FKS1* (Fig. 6A-B). Interestingly, expression of both *FKS2* (Fig. 6A) and *FKS1* (Fig. 6B) was significantly decreased in hyperactive *TOR1*^*L2134M*^ cells, following treatment with Tm. Given that Ca^2+^/ calcineurin and CWI signaling converge to mediate *FKS1/2* expression (70, 72), we differentially assessed the activity of these pathways. There was no evidence that the Ca^2+^/ calcineurin pathway was impaired in presence of Tm-induced ER stress (Supplementary Fig. 1). Additionally, we examined the activation of Rlm1 – another transcription factor regulating cell wall integrity– by assessing the expression of its downstream target, *PRM5* (Fig. 6C). We found that activation of the Rlm1 branch was not impaired in hyperactive *TOR1*^*L2134M*^ cells (Fig. 6C). Taken together, our results support the notion that defects in the cell wall architecture of hyperactive *TOR1*^*L2134M*^ mutants may be due to dysregulation of other regulators of the cell wall integrity such as the SWI4/6-SBF complex. More comprehensive studies will be required to uncover the complex role of TORC1 in the control of cell wall biogenesis and maintenance.

**Figure 6:**
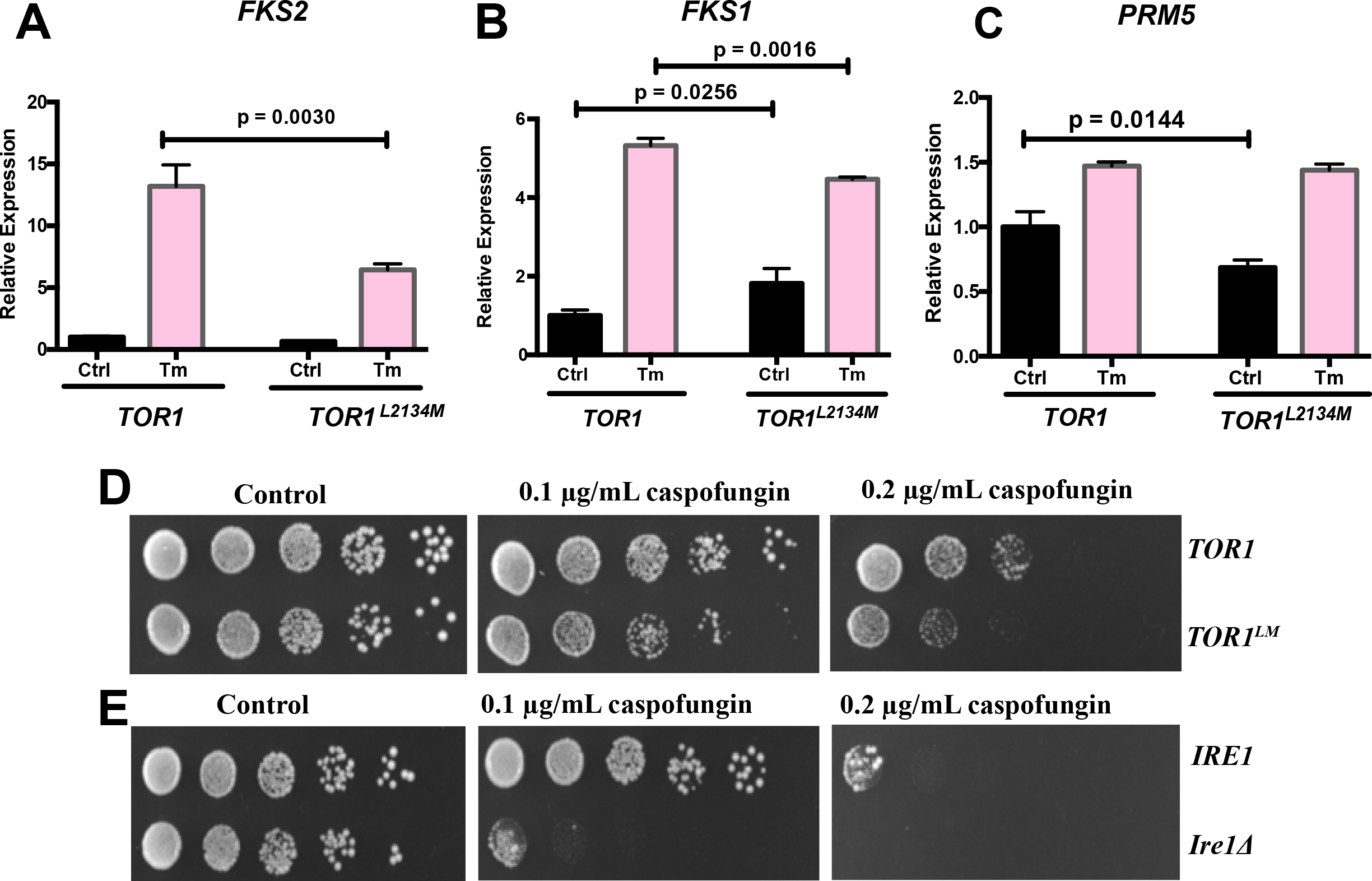
Cell wall perturbations in hyperactive *TOR1*^*L2134M*^ cells may be due to defects in glucan synthesis. **(A)** Cells expressing *WT TOR1* or hyperactive *TOR1*^*L2134M*^ were treated with tunicamycin (Tm; 2.5 μg/mL) for 2 hrs. Tm induced a significant decrease in the expression of glucan synthase genes *FKS2* and **(B)** *FKS1* as measured by qRT-PCR (n=3; ± SD). **(C)** qRT-PCR was also used to assess the expression of the Rlm1 target, *PRM5* (n=3; ± SD). **(D)** Cell growth of WT *TOR1* and *TOR1*^*L2134M*^ cells was assessed by serial dilutions on YPD plates supplemented with various concentrations of caspofungin. Compared to WT TOR1, hyperactive *TOR1*^*L2134M*^ cells displayed reduced growth. **(E)** Growth of wild-type cells and *Ire1*Δ cells was assessed by serial dilutions on YPD plates supplemented with various concentrations of caspofungin.

Given that the cell wall is essential for fungal survival and its composition is unique to the fungal organism, this structure acts as an ideal target for antifungal drugs (73). Notably, echinocandins represent the first class of antifungal drugs that specifically target the fungal cell wall (74, 75). In particular, the echinocandin caspofungin acts as a fungicide by noncompetitively inhibiting the β-1-3-glucan synthases, Fks1 and Fks2, thereby blocking cell wall synthesis (76). Since our results indicate that hyperactive *TOR1*^*L2134M*^ impairs *FKS2* and *FKS1* synthesis, we investigated whether this defect sensitizes cells to the antifungal, caspofungin (Fig. 6D). Indeed, cells expressing hyperactive *TOR1*^*L2134M*^ exhibited a growth defect as compared to WT *TOR1* cells, and this defect was further exacerbated with increasing concentrations of caspofungin (Fig. 6D). To further elucidate the connection between ER stress signaling and sensitivity to antifungal drugs, we examined the growth of *ire1Δ* cells following treatment with caspofungin (Fig. 6E). Compared to wild-type strains, *ire1Δ* showed hypersensitivity to caspofungin, suggesting that a functional ER stress response is required for resistance to this antifungal drug (Fig. 6E). Similarly, UPR-deficient strains of pathological fungi such as *C. neoformans* and *A. fumigatus* show decreased virulence in animal models (77–80). Interestingly, deletion of *MDS3* in *Candida albicans* leads to TORC1 hyperactivation resulting in filamentation defects, supporting a negative role for TORC1 hyperactivation in pathogenicity (81). Conversely, reduced TORC1 signaling in *oma1*Δ strains resulted in attenuated TORC1 signaling and increased virulence in *Candida albicans* (82). Thus, the amplitude of TORC1 signaling emerges as an important determinant of the capacity of *C. albicans* cells to withstand stress such as oxidative stress (83) and perhaps ER stress, thus impacting its virulence and pathogenicity.

While initially described as distinct pathways, our research points to a functional interaction between the UPR, TORC1, and CWI signaling pathways. Here, we use a hyperactive variant of *TOR1* to present a novel mechanism of ER stress regulation by TORC1 signaling. We show that attenuation of TORC1 signaling is required for adaptation to ER stress, and that hyperactive TORC1 signaling results in compromised cell wall architecture. Taken together, we propose that hyperactivation of TORC1 signaling alters cell wall composition, sensitizing cells to ER stress causing agents such as antifungal drugs.

## Conclusion

The high prevalence of pathogenic fungal infections, coupled with the emergence of new fungal pathogens, has rapidly brought these diseases to the forefront of global health problem. Of particular concern are the millions of people worldwide that will contract life-threating invasive fungal infections (IFI) – diseases with a mortality rate which exceeds 50%, even with the availability of antifungal treatments (84, 85). As a whole, the aetiological agents responsible for more than 90% of IFI-related deaths fall largely within four genera of fungi: *Cryptococcus, Candida, Aspergillus*, and *Pneumocytis* (84, 86). While antifungal treatments have advanced over the last decade, patient outcomes have not substantially improved (87). These shortcomings are largely attributed to the evolutionary similarity between fungi and humans, which limits the scope of drug development against fungal specific targets. As such, there is a pressing need to understand the unique cellular mechanisms that govern fungal viability. Given that *S. cerevisiae* is evolutionarily related to a number of pathogenic fungi, and in particular to the *Candida* species (88), most genes from *S. cerevisiae* are highly conserved in pathogenic fungal strains. Among the shared genomic features includes similar mechanisms for cell wall homeostasis (89–91) and activation of stress responses (92). Here we show that hyperactivation of TORC1 signaling sensitizes yeast cells to both ER stress and cell wall stressors by compromising cell wall integrity. Therefore, targeting TORC1 signaling and ER stress pathways may be useful in developing novel targets for antifungal drugs.

## MATERIALS AND METHODS

### Yeast strains and methods

The *Saccharomyces cerevisiae* strains and plasmids used in this study are listed in Tables 2.1 and 2.2, respectively. All yeast strains are derivatives of BY4742. The TS161 (*TOR1*) and TS184 (*TOR1*^*L2134M*^) strains were kind gifts from Dr. Maeda (24). BY4742 or derivatives were thawed from frozen stocks and grown on YPD (yeast extract peptone dextrose) or selective SC (synthetic complete) media for 2 days at 30°C before being transferred to liquid cultures. All experiments were carried out using either SC media containing 2% wv^−1^ glucose supplemented with 100× inositol or YPD media. Cultures were grown at 30°C with constant agitation or on selective agar plates.

**Table 1:** Yeast Strains

| Strains | Genotype | Reference |
| --- | --- | --- |
| BY4742 | *MATα his3Δ1 leu2Δ0 lys2Δ0 ura3Δ0* | (93, 94) |
| TS161 | *MATα ura3-52* | (24) |
| TS184 | *MATα ura3-52 TOR1L2134M* | (24) |
| BY4742 *ire1Δ* | *MATα his3Δ1 leu2Δ0 lys2Δ0 ura3Δ0 IRE1::KAN* | Deletion collection |

**Table 2:** Plasmids

| Plasmids | Number | Vector Backbone | Resistance | Reference |
| --- | --- | --- | --- | --- |
| pPM47 (UPR-RFP CEN/ARS URA3) | Addgene plasmid # 20132 | pRS316 | URA | (45) |
| pAMS366 (4X *CDRE-lacZ* URA3) | – | pAMS366 | URA | (95) |
| pRS316 BCK1-20 | – | pRS316 | URA | (96) |
| pRS416 GPD | ATCC 87360 | pRS416 | URA | (97) |

### Spotting and liquid growth assays

Cell growth was assessed by both spot assay and liquid culture as previously described by Duennwald (2013). Briefly, spotting assays were performed with yeast cells that were cultured overnight in selective media with 2% glucose as the sole carbon source. Cells were then diluted to equivalent concentrations of OD_600_ 0.2 and were spotted in 4 sequential five-fold dilutions. Equal spotting was controlled by simultaneously spotting cells using a multi-channel ultra-high-performance pipette (VWR International). Cells were grown on selective plates at 30°C for 2 days and imaged using a Geldoc system (Bio-RAD). For liquid cultures cells were diluted to OD_600_ 0.15 and incubated at 30°C. OD_600_ was measured every 15 mins using a BioscreenC plate reader (Growth curves USA) for 24 h. Growth curves were generated and the area under the curve was calculated for biological replicates. Statistical significance was determined using a two-tailed student T-test and GraphPad (Prism).

### Yeast Transformation

Yeast transformations were performed using the lithium acetate transformation protocol as previously described(98). Briefly, 1 mL of OD_600_ = 1, overnight cultures were pelleted at 3000 xg for 1 min. Cells were aspirated and washed with 1.5 mL sterile 0.1 M LiAc in TE buffer. Cells were then pelleted and resuspended in 285 μL sterile 50% PEG 4000 in 0.1M LiAc, 2.5 μL plasmid, and 10 μL boiled salmon sperm DNA, and incubated at 30°C for 45 mins. After that, 43 μL of sterile DMSO was added and cells were heat shocked for 15 min at 42°C before being plated on amino acid selection plates.

### Drugs

Stock solutions of tunicamycin (5 μg mL^−1^ in DMSO; Amresco), calcofluor white (30 mg mL^−1^ in H_2_O; Sigma Aldrich), rapamycin (1 mg ml^−1^ in DMSO; Fisher Bioreagents), sorbitol (3 M in H_2_O; Fisher Bioreagents), and fluorescent brightener 28 (Calcofluor white stain; 25μM; Sigma Aldrich) were used at the indicated concentrations.

### Stress Condition Experiments

In all the experiments, yeast cultures were grown to log phase (OD_600_ ~0.3) before being exposed to different stress conditions. Endoplasmic reticulum stress was achieved by adding 0.5 μg mL^−1^, 1.0 μg mL^−1^, or 2.5 μg mL^−1^ tunicamycin (Amresco) or by inositol withdrawal. For inositol depletion experiments, cells were washed twice in SC media (YNB-Inositol; Sunrise Science) and then resuspended into pre-warmed SC media lacking inositol. Cell wall stress was achieved by adding 5-20 μg mL^−1^ calcofluor white. Sorbitol rescue assays were facilitated by adding 1 M sorbitol to the media.

### qRT-PCR

RNA extraction was performed using the MasterPure Yeast RNA Purification Kit (Epicentre). cDNA was synthesized using the RevertAid H Minus First Strand cDNA Synthesis Kit (Thermoscientific). The cDNA preparations were used as templates for amplification using SsoAdvanced^Tm^ Universal SYBR ^®^ Green Supermix (Bio-Rad). The primers used are listed in Table 3. The relative expression levels were calculated using the comparative Ct method with *U3* as a reference gene.

**Table 3:**
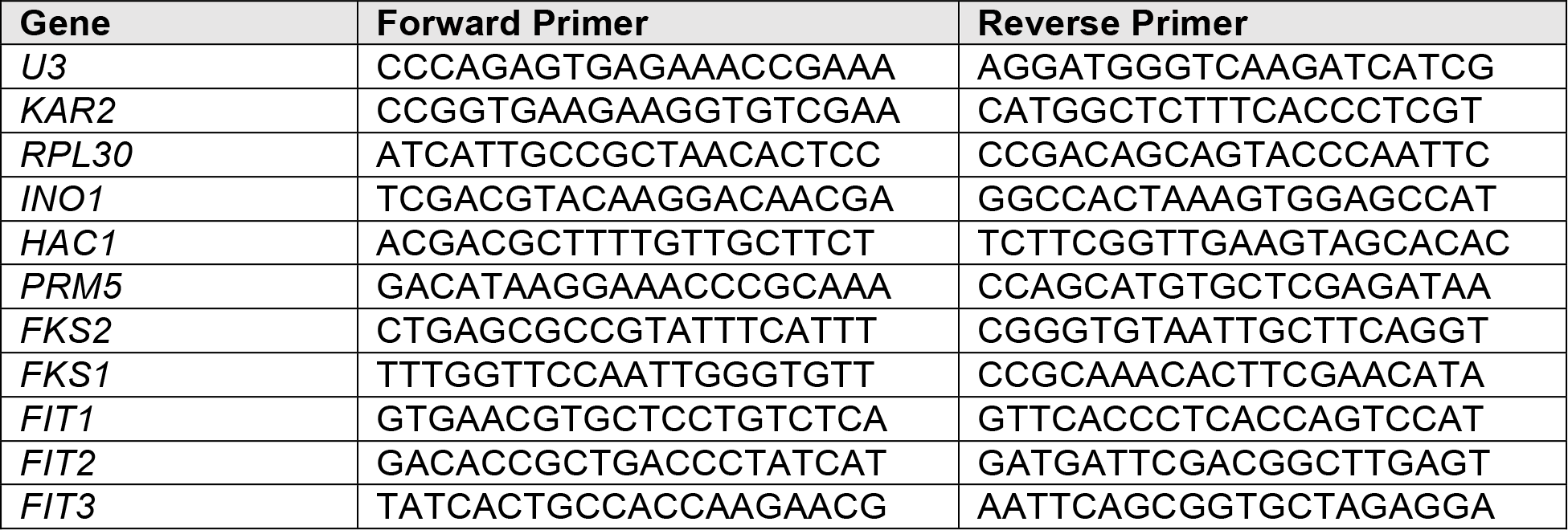
Primers

### Fluorescence Microscopy

*TOR1* and *TOR1*^*L2134M*^ cells expressing a UPR-mcherry fluorescent reporter were grown to mid-log phase before being treated with 2.5 μg mL^−1^ tunicamycin (Amresco) or inositol withdrawal for 3 h. Cells were diluted 10X, transferred to a 96 well plate, and imaged at room temperature. Fluorescence microscopy was performed using the Cytation 5 Cell Imaging Multi-Mode Reader (BioTek); the 20X objective lens and Texas Red Filter cube (586 647^−1^ nm) were used. Images were analyzed using ImageJ software (https://imagej.nih.gov/ij/). Violin plots presented in Figure 2D were generated using the PlotsOfData software (99).

### *HAC1* Splicing Assay

Cells were cultured to mid-log phase before being treated with either 1.0 μg/mL tunicamycin (Amresco) or inositol withdrawal for 2 h. RNA extraction was performed using the MasterPure Yeast RNA Purification Kit (Epicentre). cDNA was synthesized from the extracted RNA using the RevertAid H Minus First Strand cDNA Synthesis Kit (Thermoscientific). The cDNA preparations were then used as templates for RT-PCR with *HAC1* primers (listed in Table 4). The resulting reaction product was separated by electrophoresis on an agarose gel and bands were visualized using a Geldoc system (Bio-Rad).

### β-galactosidase Assay

*TOR1* and *TOR1*^*L2134M*^ yeast strains transformed with plasmids carrying the *CDRE-LacZ* reporter were assayed as previously described (100). Briefly, cells were grown to log phase in selective SC media, harvested by centrifugation, then cultured in SC media containing the indicated concentrations of stressors or CaCl_2_. After incubation at 30°C for 2 h, cells were harvested by centrifugation and resuspended in lacZ buffer. To measure β-galactosidase activity, 50 μL cell lysate was mixed with 950 μL lacZ buffer containing 2.7 μL β-mercaptoethanol, 1 drop 0.1% SDS, 2 drops CHCl_3_ and incubated at 30°C for 15 min. The reaction was started by adding 100 μL ONPG (4 mg mL^−1^) and incubated at 30°C till the colour changed to yellow. The reaction was stopped by adding 300 μL of 1 M Na_2_CO_3_. β-galactosidase activity was determined at 420 nm absorbance using a plate reader, normalizing data to cell density.

### Protein Extraction and Western Blot

Cells were lysed using alkaline lysis with 0.1 M NaOH (101) and proteins were extracted into 4x Laemmli sample buffer containing 100 mM DTT. Protein samples were separated using SDS-PAGE (BioRad Mini-PROTEAN TGX Pre-Cast gels, 4-15%) and transferred to nitrocellulose membranes using the BioRad Trans-Blot^®^ Turbo^™^ RTA Transfer Kit. Membranes were blocked with 5% fat free milk for 30 mins, before probing with P-S6 Ribosomal Protein S235 236^−1^ Rabbit Ab (Cell Signaling Technology) or anti-PGK1 (Invitrogen) overnight at 4°C. Membranes were then incubated with the Alexa Fluor 488 goat anti-rabbit for 1 hr. Membranes were imaged using a BioRad infrared imager (BioRad).

### Calcofluor White Stain Microscopy and Flow Cytometry

*TOR1* and *TOR1*^*L2134M*^ cells were grown in triplicate to mid-log phase in YPD media, before being treated with Fluorescent Brightener 28 (Sigma-Adlrich) to a final concentration of 25 μM. Cells were grown for 20 min at 30°C with continuous shaking before they were pelleted and washed in SC media. Cells were diluted 10× in growth media and plated in Lab-Tek (Thermo Inc.) imaging chambers and processed for fluorescence microscopy. Images were acquired using a Zeiss AxioVert A1 wide filed fluorescence microscopy equipped with a 63X NA 1.4 Plan Apopchromat objective, 359 nm excitation 461 nm^−1^ emission (DAPI) long pass filter and an AxioCam ICm1 R1 CCD camera (Carl Zeiss inc.). Images were analyzed using ImageJ software. For flow cytometric analysis, cells were cultured in appropriate media and processed for flow cytometry using a BD Bioscience FACS Celesta flow cytometer equipped with a 405 nm Violet laser. Data was analyzed using the BD FACS Diva Software. All conditions were performed in triplicate, 20 000 cells were analyzed, and mean fluorescence intensities were calculated. No gates were applied.

### Microarray Analysis

*TOR1* and *TOR1*^*L2134M*^ yeast cultures were grown to log phase (OD_600_ ~0.3) before being treated with tunicamycin (2.5 μg/mL). RNA was extracted from two independent cultures (n=2) and quality was assessed with Bioanalyzer as previously described (102). Microarray analysis was conducted with the GeneChip^®^ Yeast Genome 2.0 Array (Affymetrix, Santa Clara, California, USA). Briefly, biotinylated complimentary RNA (cRNA) was prepared from 100 ng of total RNA as per the GeneChip 3’ IVT PLUS Reagent Kit manual (ThermoFisher Scientific, Waltham, MA). (https://www.thermofisher.com/order/catalog/product/902416). Data was analyzed using the Transcriptome Analysis Console (TAC) software (Affymetrix) by filtering for genes that showed a two-fold change in expression with a p-value of 0.05 using sacCer3 as a reference genome. Gene lists were created using the gene ontology term finder on the *Saccharomyces* genome database (https://www.yeastgenome.org/). All microarray data were submitted to the GEO database as series GSE129200.

## ACKNOWLEDGMENTS

We thank Dr. Tasuya Maeda (University of Tokyo), Dr. Martha Cyert (Stanford University), and Marina Molina (Universidad Complutence Madrid) for the yeast plasmids and strains. This work is supported by a National Science and Engineering Research Council of Canada (NSERC) discovery grant (RGPIN-2015-06400) to PL and an NSERC Canada Graduate Scholarship to KA. PL is the recipient of a John R. Evans Leaders Fund award (#35183) from the Canadian Foundation for Innovation (CFI) with matching fund from the Ontario Research Fund. The authors thank Dr Christopher Brandl, Dr Martin Duennwald, and Dr Rebecca Shapiro for useful discussions about the manuscript.

---

**Supplemental Figure 1:**
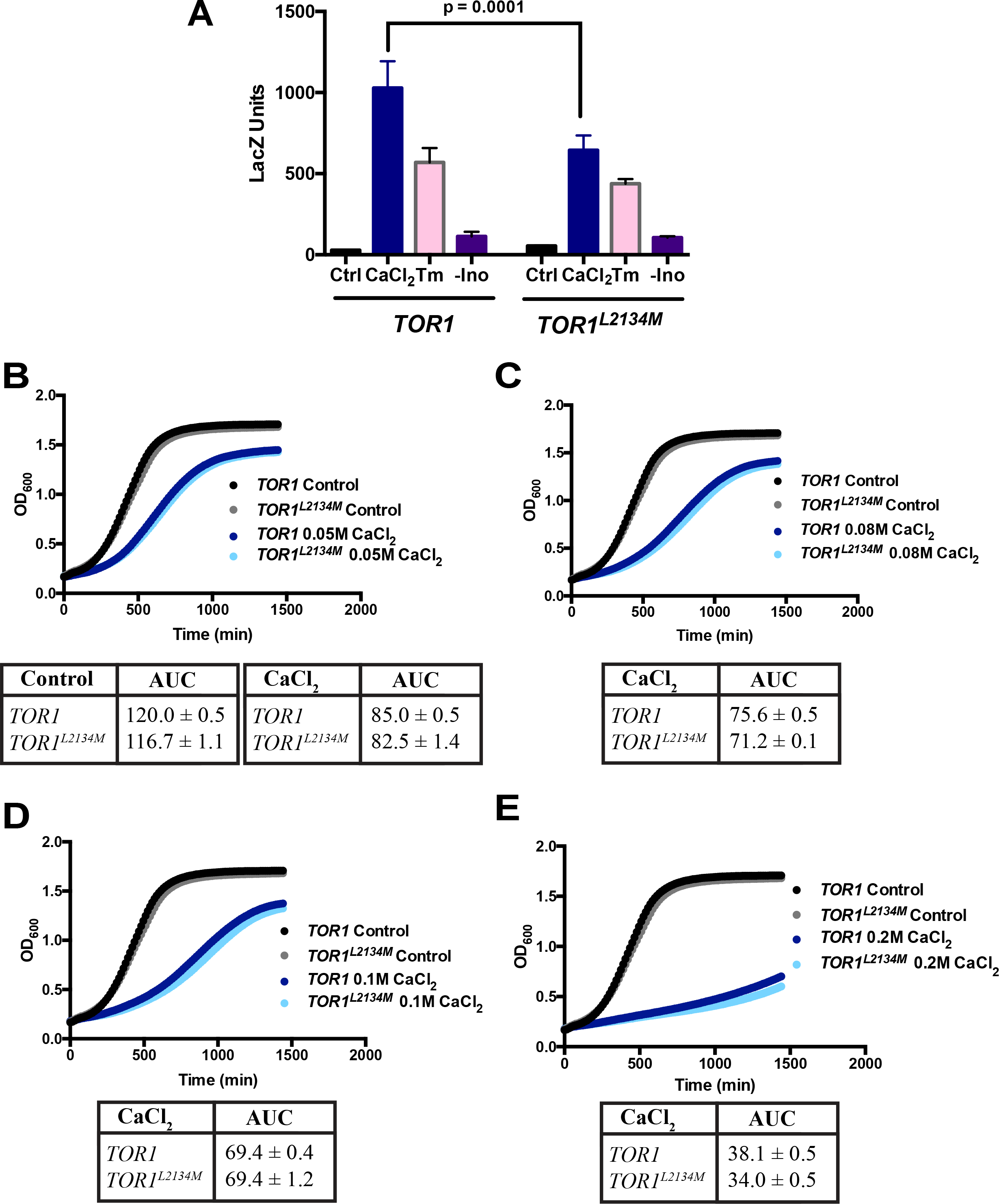
The Ca^2+^/calcineurin pathway is not impaired in hyperactive *TOR1*^*L2134M*^ mutants. **(A)** β-galactosidase activity (measured in LacZ units) was used to assess expression of calcineurin dependent response element (CDRE) following treatment with CaCl_2_ (1 M), tunicamycin (Tm; 1.0 μg/mL), or inositol withdrawal (-ino; n=6). **(B-E)** Growth of cells expressing WT *TOR1* or hyperactive *TOR1*^*L2134M*^ was assessed by liquid growth assay following treatment with 0.05 M CaCl_2_, 0.08 M CaCl_2_, 0.1 M CaCl_2_, or 0.2 M CaCl_2._ The area under the curve (AUC) was quantified for each replicate (n=3). All conditions were run simultaneously. Control conditions are reproduced on each panels for clarity.

